# Mechano-mediated M2 macrophage polarization and immune suppression in stiffened tumor microenvironment

**DOI:** 10.1101/2024.07.29.605566

**Authors:** Jiranuwat Sapudom, Paul Tipay, Jeremy CM Teo

## Abstract

The tumor microenvironment (TME), which is composed of various cell types and the extracellular matrix (ECM), plays crucial roles in cancer progression and treatment outcomes. However, the impact of the mechanical properties of the ECM, specifically collagen fibril alignment and crosslinking, on macrophage behavior and polarization is less understood. To investigate this, we reconstituted 3D collagen matrices to mimic the physical characteristics of the TME. Our results demonstrated that stiffening the matrix through the alignment or crosslinking of collagen fibrils promotes macrophage polarization toward the anti-inflammatory M2 phenotype. This phenotype is characterized by increased expression of CD105 and CD206 and a distinct cytokine secretion profile. The increased stiffness and aligned fibrils activate mechanotransduction pathways, notably integrin β1 and PI3K signaling, leading to increased IL-4 secretion, which acts in an autocrine manner to further promote M2 polarization. Interestingly, these stiffened microenvironments also suppressed the proinflammatory response. In coculture experiments with breast cancer cell lines (MDA-MB-231 and MCF-7), macrophages within stiffened or aligned matrices significantly increased cancer cell proliferation and invasion. These findings suggest that the mechanical properties of the ECM, specifically its alignment and crosslinking, create a more favorable environment for tumor progression by modulating macrophage activity. Overall, our study underscores the critical role of ECM mechanics in shaping immune cell behavior within the TME, highlighting the potential for therapies that target ECM properties and macrophage polarization to inhibit cancer progression and enhance treatment efficacy.

## 1. Introduction

The complex relationship between cancer progression and treatment efficacy is intricately tied to the tumor microenvironment (TME), a dynamic cellular environment encompassing diverse cell types, the extracellular matrix (ECM), and various biochemical mediators [1–4]. Within this microenvironment, the ECM extends beyond its structural role, significantly influencing cellular behavior, orchestrating cell signaling, and governing crucial functions such as cell proliferation, motility, and differentiation [5]. As cancer progresses, the TME undergoes significant physical and biochemical changes, including excessive production of ECM components by cancer-associated fibroblasts (CAFs) [6, 7] and ECM stiffening due to increased enzymatic cross-linking by lysyl oxidase and enhanced collagen fibril alignment [8–10]. These alterations collectively create a distinct mechanical microenvironment that impacts the behavior of cancer cells and stromal cells, especially immune cells and fibroblasts, by modulating their migration and activation within the TME [11, 12]. For example, a stiffened ECM activates mechanotransduction pathways (e.g., the YAP/TAZ, Rho/ROCK, and cell nuclear deformation pathways), promoting cancer cell survival, proliferation, migration, and drug response [2, 13, 14]. Additionally, the reorganization of ECM fibrils into aligned structures and microarchitecture facilitate fibroblast-to-myofibroblast differentiation via cell contractility [15, 16], reduces the T-cell immune response via YAP signaling [17], and enhances cancer cell invasion, potentially aiding metastasis [18–20]. Understanding these ECM changes and deciphering their functions in regulating cancer and stromal cells is essential for developing novel and improved therapies [21, 22].

Macrophages are known to influence the balance between the inhibition and promotion of cancer cells during cancer development and metastasis [23, 24]. They exhibit remarkable plasticity, allowing them to change their physiology in response to various environmental stimuli. While macrophages exist across a spectrum of activation states, they can traditionally be classified into two main types: proinflammatory (M1) and anti-inflammatory (M2) macrophages [25]. M1 macrophages are generally associated with tumor-suppressing functions, including promoting inflammation, producing reactive oxygen species, and facilitating antigen presentation to T cells. These activities help attack and destroy cancer cells in the early stages of tumor development [13, 24]. In contrast, M2 macrophages are associated with tissue repair and immunosuppression, facilitating tumor growth and metastasis. M2 macrophages can be further divided into distinct subtypes, such as M2a, M2b, M2c, and M2d, based on marker expression and cytokine secretion profiles [25, 26]. These subtypes perform various functions, including wound healing, immune regulation, and angiogenesis support, which can create a tumor-promoting environment [26]. An increased M1/M2 macrophage ratio is correlated with enhanced survival rates and improved clinical outcomes, as the presence of more M1 macrophages can lead to a more robust antitumor immune response [23, 27]. In the TME, macrophages expressing CD206, also known as the mannose receptor, are often found within and adjacent to cancer tissues. These CD206-expressing macrophages tend to accumulate in cancer tissues in mouse models [28] and human biopsies [29–31]. These macrophages, often classified within the M2 subspectrum, specifically M2a macrophages, are associated with triggering fibroblast differentiation into myofibroblasts via TGF-β1 secretion [32, 33]. Myofibroblasts are essential for the formation of the ECM, providing structural support to tumors. The ECM supports tumor cell adhesion and migration and contributes to tumor tissue stiffness, which is often linked to a more aggressive tumor phenotype and poor prognosis [6]. Additionally, M2a macrophages are known for their immunosuppressive properties [34]. By secreting anti-inflammatory cytokines such as IL-10, they help create an immunosuppressive microenvironment [32]. This suppression of the immune response allows tumor cells to evade immune surveillance, enabling unchecked tumor growth and spread [27, 33, 35, 36].

Recent studies have attempted to explore the impact of the mechanical attributes of the ECM on macrophage phenotypes and functions, mostly using mechanically adjustable two-dimensional cell culture models or three-dimensional hydrogels that lack the microarchitectural and fibrous features of native tissues [37, 38]. To address this limitation, three-dimensional (3D) collagen-based matrices can be reconstituted and independently fine-tuned for various biophysical and biochemical properties, including porosity [39], fibril characteristics [16, 40], crosslinking types and degrees [41], bulk and fibril bending stiffness [42], and the incorporation of other ECM components [43, 44].

In this work, we reconstituted 3D collagen-based matrices to replicate distinct biophysical features of the tumor microenvironment (TME), such as collagen fibril alignment and crosslinking. We aimed to demonstrate how specific ECM characteristics, such as microarchitecture (random vs. aligned fibrils) and mechanical properties, modulate macrophage phenotypes and functions. We also investigated whether these ECM characteristics suppress macrophage inflammatory responses and explored the underlying mechanotransduction mechanisms. Additionally, we co-cultured MDA-MB-231 and MCF7 breast cancer cell lines within our 3D collagen matrices to observe the direct impact of ECM characteristics on cancer cell behavior and macrophage‒cancer cell interactions. We assessed how ECM features influence cancer cell proliferation, migration, invasion, and macrophage polarization. Our findings could reveal novel mechanisms by which the biophysical features of the ECM within the TME help cancer cells evade immune defenses, potentially leading to innovative cancer treatments.

## 2. Materials and methods

### 2.1 Reconstitution of collagen matrices

To reconstitute 3D collagen matrices, a collagen solution was prepared using an established protocol [45]. Briefly, rat tail type I collagen (Advanced BioMatrix, Inc., Carlsbad, CA, USA) was mixed with 0.1% acetic acid (Merck KGaA, Darmstadt Germany) and 0.5 M phosphate buffer (Merck KGaA, Darmstadt, Germany) to achieve a final collagen concentration of 2 mg/mL. The prepared collagen solution was then transferred onto glutaraldehyde-coated coverslips (13 mm in diameter; VWR, Darmstadt, Germany) and placed on a planar surface for a random matrix or on a custom-designed inclined surface with an inclination angle of 30° for an aligned matrix, as previously established [15]. Collagen fibrillogenesis was initiated at 37°C, 5% CO2, and 95% humidity. The reconstituted 3D collagen matrices were then washed three times with phosphate-buffered saline (PBS; Sigma‒Aldrich, Schnelldorf, Germany) and kept in PBS prior to further experiments.

To crosslink the reconstituted collagen matrices, the matrices were incubated with 4 mg/mL N-(3- dimethylaminopropyl)-N′-ethylcarbodiimide hydrochloride (EDC; Sigma‒Aldrich, Schnelldorf, Germany) dissolved in 0.1 M 2-(N-morpholino)ethanesulfonic acid (MES; Sigma‒Aldrich, Schnelldorf, Germany) at pH 5 for 2 hours at room temperature, following previously published methods [42]. Following crosslinking, the matrices were washed three times with PBS (Thermo Fisher Scientific Inc., Leicestershire, UK), incubated with cell culture medium for 24 hours, and washed three times with PBS before use.

### 2.2 Characterization of the topological and mechanical properties of reconstituted 3D matrices

For the analysis of topological parameters, reconstituted collagen matrices were stained with 50 µM 5-(and 6)-carboxytetramethylrhodamine succinimidyl ester (TAMRA-SE; Sigma‒Aldrich, Schnelldorf, Germany) at room temperature for 60 minutes and then washed three times with PBS. The matrices were subsequently visualized using a confocal laser scanning microscope (cLSM; SP8 Leica, Wetzlar, Germany) with a 40× water immersion objective (Leica, Wetzlar, Germany). To quantify the topological properties of the 3D collagen matrices, the acquired images were analyzed using a custom-built MATLAB script (MATLAB 2024b; MathWorks, Portola Valley, CA, USA), as described in a previous publication [46]. Using stacked images obtained from four different random positions per matrix, the pore diameter, fibril diameter, and degree of fibril alignment of the reconstituted collagen matrices were analyzed.

To measure the mechanical properties of the reconstituted matrices, a nondestructive rheological method using ElastoSens™ Bio (Rheolution, Quebec, Canada) was used, as previously published [15]. All the experiments were performed in four replicates.

### 2.3. Macrophage differentiation

The human monocytic THP-1 cell line (AddexBio, San Diego, CA, USA) was maintained in RPMI- 1640 cell culture medium supplemented with 10% fetal bovine serum (FBS), 1% penicillin/streptomycin, 1% 4-(2-hydroxyethyl)-1-piperazineethanesulfonic acid (HEPES), 1% sodium pyruvate, and 0.01% beta-mercaptoethanol under standard cell culture conditions of 37°C, 5% CO2, and 95% humidity. All cell culture reagents were purchased from Thermo Fisher Scientific, Inc., Leicestershire, UK.

For the cell studies, 2 × 10^5^ THP-1 cells were seeded onto prepared collagen matrices in FBS-free RPMI-1640 cell culture medium and placed in an incubator overnight to allow infiltration of the cells into the collagen matrices. To induce differentiation into uncommitted macrophages (M0), cells were cultured under standard cell culture conditions in FBS-free RPMI-1640 cell culture medium supplemented with 300 nM phorbol 12-myristate 13-acetate (PMA; Merck KGaA, Darmstadt, Germany) for 6 hours, as previously published [39]. Afterwards, the differentiation medium was removed, and the cell-embedded matrices were washed three times with PBS and incubated for 24 hours in FBS-free RPMI-1640 cell culture medium prior to further investigation.

### 2.4. Blocking experiment

After uncommitted macrophages (M0) were seeded onto 3D collagen matrices, the cells were treated with mechanosensing and transduction inhibitors using an antibody against integrin β1 (2 µg/mL; anti-integrin β1; Biolegend, USA) and chemical inhibitors against PI3K (1 µM; LY294002), ROCK (1 µM; Y-27632), YAP (1 µM; verteporfin), and Piezo1 (5 µM; GsMTx4) signaling in RPMI-1640 cell culture medium. All chemical inhibitors were purchased from TOCRIS (Minneapolis, MN, USA). The cells were cultured for 48 hours under standard culture conditions. The cells were subsequently analyzed for CD206 expression and IL-4 production using a flow cytometer. The experiment was conducted in four independent replicates.

### 2.5. Quantitative analysis of macrophage surface markers and viability using flow cytometry

To evaluate the expression of cell surface markers, the matrices were digested with 10 U/mL Liberase™ DL Research Grade (Sigma‒Aldrich, Schnelldorf, Germany) for 20 minutes under standard cell culture conditions. The prepared Liberase solution was supplemented with Human TruStain FcX™ Fc receptor blocking solution at a 1:500 dilution. The cells were subsequently stained with mouse anti-human antibodies against CD105, CD163, CD206, HLA-DR, and Zombie Violet (a viability dye) at a 1:1000 dilution in PBS on ice for 30 minutes. All the antibodies were diluted in PBS at a 1:500 ratio. These staining antibodies and the viability dye were purchased from Biolegend (San Diego, CA, USA). Details of all the antibodies used are provided in Supplementary Table S1. The stained cells were analyzed using an Attune NxT flow cytometer equipped with an autosampler (Thermo Fisher Scientific, CA, USA). Data analysis was performed using FlowJo software (Becton, Dickinson and Company, NJ, USA). The experiments were conducted in 6 replicates.

### 2.6. Quantitative analysis of cytokine secretion profiles

Cell culture supernatants were collected after 3 days of culture. The LEGENDplex™ Human Essential Immune Response Panel (13-plex, including IL-4, IL-2, CXCL10 (IP-10), IL-1β, TNF-α, CCL2 (MCP-1), IL-17A, IL-6, IL-10, IFN-γ, IL-12p70, CXCL8 (IL-8), and free active TGF-β1) (BioLegend, San Diego, CA, USA) was used to quantify the secreted cytokines in the cell culture supernatant following the manufacturer’s instructions. The samples were analyzed using an Attune NxT flow cytometer equipped with an autosampler (Thermo Fisher Scientific Inc., Leicestershire, UK). Quantitative analysis was subsequently performed using the LEGENDplex™ Data Analysis Software Suite (VigeneTech, Carlisle, MA, USA). The experiments were conducted in 6 replicates.

### 2.7. RNA isolation, sequencing and functional transcriptome analysis

RNA was extracted using TRIzol (Thermo Fisher Scientific Inc., Leicestershire, UK) and chloroform (Sigma‒Aldrich, Schnelldorf, Germany) following the manufacturers’ protocols. The samples were then purified using an RNeasy Mini Kit (Qiagen, Hilden, Germany) according to the provided instructions. The RNA concentration and purity (ratio of absorbance at 260 and 280 nm) were measured with a Nanodrop (Thermo Fisher Scientific Inc., Leicestershire, UK) and confirmed using a Qi RNA kit with a Qubit 4 fluorometer (Thermo Fisher Scientific Inc., Leicestershire, UK) prior to RNA sequencing.

Purified RNA samples were prepared with an NEB Ultra II RNA kit (New England Biolabs, Ipswich, MA, USA) following the manufacturer’s instructions, utilizing the NEBNext Poly(A) mRNA Magnetic Isolation Module (New England Biolabs, Ipswich, MA, USA) for unique dual indexing. The library concentration, size distribution, and quality were assessed using a Qubit 4 fluorometer (Thermo Fisher Scientific Inc., Leicestershire, UK) with a dsDNA high-sensitivity kit (Thermo Fisher Scientific Inc., Leicestershire, UK) and a 4200 TapeStation with a High Sensitivity D5000 kit (Agilent, Santa Clara, CA, USA). Based on these results, libraries were normalized by molarity, pooled, and quantified using a library quantification kit for Illumina platforms (Roche, Basel, Switzerland) on a StepOnePlus qPCR machine (Thermo Fisher Scientific Inc., Leicestershire, UK). The pooled libraries were then loaded at 350 pM with 1% PhiX on an S2 FlowCell and subjected to paired-end sequencing (2 × 150 bp) on a NovaSeq 6000 sequencer (Illumina, San Diego, CA, USA). The RNA-Seq experiments were performed in triplicate, and the resulting RNA sequencing data were processed before transcriptome analysis, as detailed in the supplementary information.

For data analysis, we used iDEP, a publicly accessible web-based software provided by South Dakota State University [47], http://bioinformatics.sdstate.edu/idep/, accessed on May 20, 2024. Differentially expressed genes (DEGs) were identified using DESeq2, with a false discovery rate (FDR) cutoff of ≤ 0.05 and a fold change (FC) of ≥ 2.0. Additionally, parametric gene set enrichment analysis (PGSEA) was conducted to predict enrichment in biological activities and pathway interactions, aiming to identify unbiased differential gene set enrichment in biological activity and curated pathway interaction databases.

### 2.8. Coculture of macrophages and cancer cells

MDA-MB-231 and MCF-7 breast cancer cells, both of which were acquired from the American Type Culture Collection (ATCC, Manassas, VA, USA), were cultured in Dulbecco’s modified Eagle’s medium (DMEM) supplemented with 10% fetal bovine serum (FBS) and 1% penicillin/streptomycin. Cultures were maintained at 37°C with 5% CO2 and 95% humidity. All cell culture reagents were purchased from Thermo Fisher Scientific, Inc., Leicestershire, UK. Prior to performing the coculture experiment, both cancer cell lines were stained with 1 µM CFSE (BioLegend, San Diego, California, USA) in FBS-free DMEM for 30 minutes under standard cell culture conditions. The staining allows distinguishing between cancer cells and macrophages via flow cytometry and imaging.

For both monoculture and coculture with macrophages, 1 × 10^5^ CFSE-stained MDA-MB-231 or MCF-7 cells were seeded onto collagen matrices or matrices embedded with M0 macrophages. The cells were then cultured in FBS-free RPMI-1640 cell culture medium under standard conditions for 3 days.

### 2.9. Quantitative analysis of cancer cell proliferation

CFSE-stained MDA-MB-231 or MCF-7 cells were seeded onto collagen matrices or matrices embedded with M0 macrophages. To quantify the number of cells, collagen was digested with 10 U/mL Liberase™ DL Research Grade (Sigma‒Aldrich, Schnelldorf, Germany) for 20 minutes under standard cell culture conditions. The cell number, an indicator of cell proliferation, was analyzed using an Attune NxT flow cytometer equipped with an autosampler (Thermo Fisher Scientific Inc., Leicestershire, UK). The experiments were performed in 6 independent replicates.

### 2.10. Imaging and quantification of cancer cell infiltration into 3D collagen matrices

Cancer cell infiltration into collagen matrices in both monocultures and cocultures with macrophages was quantitatively determined by analyzing the nuclei of individual CFSE-positive cells. Z-stack images with a spacing of 5 µm were obtained using an epifluorescence microscope with an automatic scanning stage (DMi8 S; Leica, Wetzlar, Germany) equipped with a ×10 dry objective (Leica, Wetzlar, Germany). The z-position of the cell nuclei as a function of migration distance was examined using a custom-built MATLAB script (MATLAB 2024b; MathWorks, Portola Valley, CA, USA), as described previously [45]. The cells located more than 20 µm below the collagen matrix surface were classified and counted as infiltrated cells. The percentage of cancer cells that migrated was determined as a percentage of the total number of cancer cells per field. Four random positions were analyzed per matrix condition in three independent experiments.

### 2.11. Statistical analysis

All experiments were conducted with a minimum of four independent replicates, unless otherwise specified. The error bars represent the standard deviation (SD). Statistical significance was determined using one-way ANOVA or two-way ANOVA followed by Tukey’s post hoc analysis, which was performed with GraphPad Prism 10 software (GraphPad Software, La Jolla, CA, USA). The significance levels were set as follows: *p < 0.05, **p < 0.01, ***p < 0.001, ****p < 0.0001.

## 3. Results and discussion

The complexity of cancer tissue raises questions as to which matrix parameters most significantly contribute to the accumulation or promotion of M2-like macrophage phenotypes, as matrix stiffening can result from various factors, including enhanced crosslinking and collagen fibril alignment [8–10]. To investigate these questions, we utilized 3D collagen matrices with increased stiffness achieved through chemical crosslinking and collagen fibril alignment. Within these matrices, we differentiated monocytic THP-1 cells into uncommitted macrophages (M0) and performed immunophenotyping, cytokine secretion profiling, transcriptome analysis, and inhibition assays of various signal transduction pathways to elucidate the mechanisms driving macrophage mechanosensing. We subsequently conducted coculture experiments with macrophages and two types of breast cancer cells, MDA-MB-231 and MCF-7 cells, to investigate cell‒cell interactions within our reconstructed 3D models.

### 3.1. Establishment of well-defined 3D collagen matrices

Like in tissues, chemical crosslinking and collagen fibril alignment increase the stiffness of 3D collagen matrices. For chemical crosslinking, reconstituted collagen matrices were postmodified using a zero-length EDC crosslinker, allowing enhancement of matrix stiffness while maintaining its microarchitecture [40, 42]. For the reconstruction of the aligned matrices, the collagen matrices were reconstituted on a 30° inclined surface, as previously described [15]. This method leverages the sedimentation effects caused by gravitational forces during collagen fibrillation on an inclined surface to generate an aligned microarchitecture.

Prior to using the matrices for cell culture, we visualized and characterized the matrices in terms of their matrix microarchitecture and elastic properties. As shown in **Figure 1A**, the matrices were fluorescently labeled with TAMRA-SE and visualized using a confocal laser scanning microscope. The random and crosslinked matrices exhibited randomly distributed collagen fibrils, whereas the aligned matrix displayed an anisotropic fibril orientation. To quantitatively validate our visual observations, we employed a custom-made image analysis toolbox to analyze the distribution of collagen fibril orientation and measure the coherence index. As shown in **Figure 1B**, the random and crosslinked matrices displayed a uniformly distributed fiber orientation, whereas the aligned matrices exhibited a Gaussian distribution. To further examine fibril orientation, we calculated the coherence index, a quantitative measure on a scale from 0 to 1, where 0 represents a random fibril distribution and 1 signifies perfect fibril alignment. Our data revealed that the coherence index was approximately 0.18 for random matrices, and the value remained consistent for both random and crosslinked matrices **(Figure 1C)**. In contrast, a significant increase in the coherence index was observed in the aligned matrices, confirming the successful reconstruction of a matrix with aligned fiber orientation, as visually observed (**Figure 1C**). Although the coherence index was 0.57, this is due to the stochastic nature of the collagen assembly, which cannot be compared to the perfect alignment found in printed materials. However, the coherence index of the aligned matrix closely resembled the collagen alignment observed in human fibrotic conditions and certain cancer tissues [18, 48].

**Figure 1:**
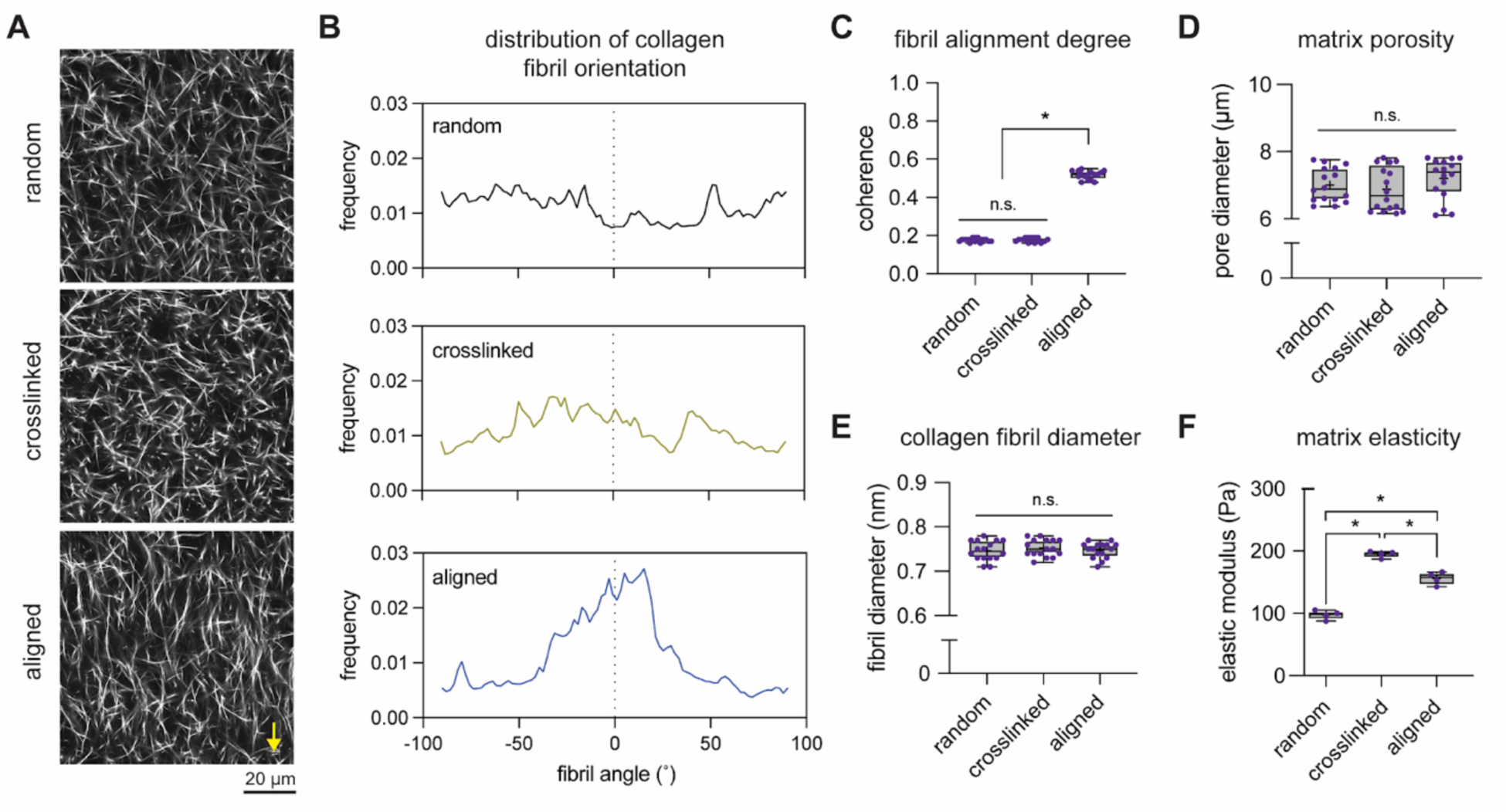
Topological and mechanical characterization of reconstituted matrices. **(A)** Representative images of collagen matrices (scale bar = 20 µm). Yellow arrows indicate the direction of fibril alignment. Matrix topology was quantified using image-based analysis toolboxes. **(B)** Representative distribution plot of the degree of collagen fibril orientation. **(C)** Quantitative analysis of the coherence index, indicating the degree of collagen fibril alignment. **(D)** Quantitative analysis of the mean pore diameter and **(E)** mean fibril diameter. For topological analysis, four different positions of each matrix condition from four different samples were analyzed. **(F)** The bulk matrix elastic modulus was quantified using a nondestructive contactless rheometer from four different samples of each matrix condition. The data are shown as box plots: the box represents the median with the 10th and 90th percentiles, the error bars indicate the minimum and maximum values, and the plus sign (+) represents the mean of the data. An asterisk (*) indicates statistical significance (p ≤ 0.05) as determined by one-way ANOVA followed by Tukey’s post hoc test.

In addition to analyzing fibril alignment, we also quantitatively characterized other topological and mechanical properties of the reconstituted matrices, including matrix porosity, collagen fibril diameter, and elasticity. Our data revealed no significant differences in matrix porosity or collagen fibril diameter among the established matrix conditions (**Figure 1D and 1E**). Quantification of the matrix elasticity revealed a significant increase in the elastic modulus of the crosslinked and aligned matrices compared with that of the random matrix (**Figure 1F**). An increase in the elastic modulus in an aligned matrix compared with a random matrix is well correlated with previous reports [15, 17, 49, 50].

### 3.2. Stiff matrices induce macrophage polarization toward the M2-like phenotype

We subsequently utilized reconstituted matrices to examine the effects of matrix stiffening on the macrophage phenotype. In our study of macrophage differentiation within these matrices, THP-1 monocytic cells were used. These cells were seeded onto the matrices and differentiated into uncommitted macrophages (M0) using PMA for 6 hours, followed by a resting period in cell culture media for 72 hours. This process of macrophage differentiation is illustrated in **Figure 2A**. Representative images of macrophages cultured on various matrices are shown in **Figure 2B**. These images revealed that macrophages consistently maintain a round morphology across the different matrices.

**Figure 2:**
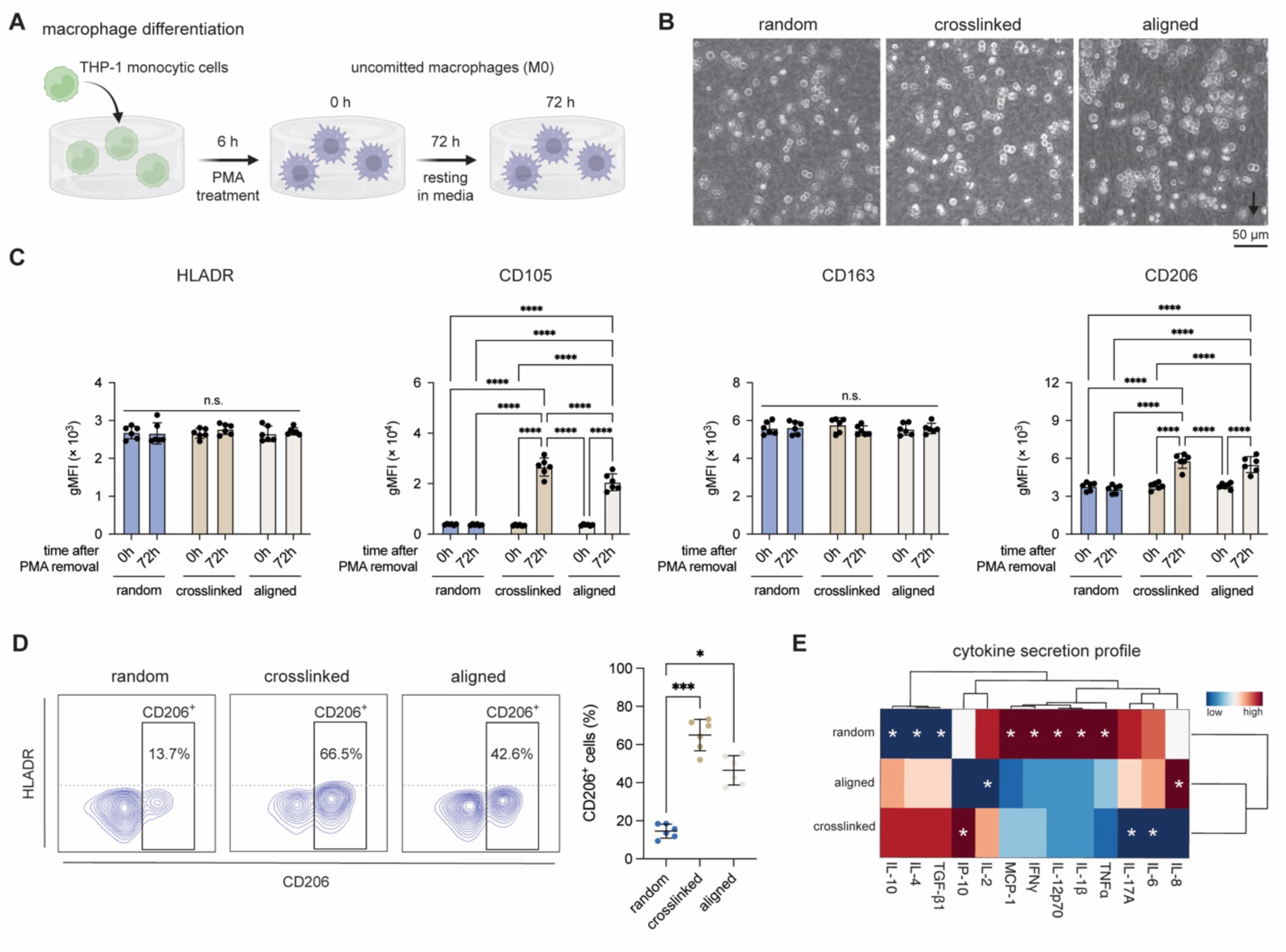
Immunophenotyping and cytokine secretion profiling of differentiated macrophages (M0) on reconstituted matrices. **(A)** A schematic illustration depicting the differentiation of macrophages within 3D collagen matrices. **(B)** Representative images displaying the morphology of differentiated macrophages after resting for 72 h on reconstituted matrices. (Scale bar = 20 µm). For immunophenotyping, the cells were stained with specific antibodies and analyzed using a flow cytometer. **(C)** Quantification of the geometric mean fluorescence intensity (gMFI) for HLADR, CD105, CD163, and CD206 in differentiated macrophages immediately after cultivation for 6 h and 72 h. **(D)** Coexpression plot of HLADR and CD105, as well as quantitative analysis of CD206-positive macrophages after 72 h of culture. **(E)** Cytokine secretion profiles were quantified using multiplex bead ELISA for macrophages after 72 h of cultivation. The data are presented as a heatmap, with blue indicating low secretion and red indicating high secretion. Levels of statistical significance were assessed using two-way ANOVA followed by Tukey’s post hoc test. Significance levels were established as follows: *p < 0.05, **p < 0.01, ***p < 0.001, ****p < 0.0001. The experiments were performed in 6 independent replicates.

To determine whether matrix stiffening influences macrophage polarization into specific phenotypes, we analyzed cell surface markers and cytokine secretion profiles. As depicted in **Figure 2C**, we measured the expression levels of HLADR, CD105, CD163, and CD206 via flow cytometry. HLADR serves as a key marker for proinflammatory (M1) macrophages, whereas CD105, CD163, and CD206 are markers for anti-inflammatory (M2) macrophages. Notably, M2a macrophages exhibit increased expression of CD105 and CD206, whereas M2c macrophages predominantly express CD163. Our results revealed increased expression of CD105 and CD206 in macrophages cultured on crosslinked and aligned matrices for 72 hours, whereas the HLADR and CD163 levels were similar to those in macrophages at 0 hours across all matrix conditions. We also examined the prevalence of CD206-positive cells, as this macrophage phenotype is enriched in the tumor microenvironment [34, 51, 52]. **Figure 2D** shows a significant increase in CD206-positive cells in both crosslinked and aligned matrices relative to random matrices, with 66.5% in crosslinked matrices and 42.6% in aligned matrices, compared with 13.7% in random matrices.

From this surface marker analysis, we inferred that matrix stiffening may facilitate a shift in the macrophage phenotype from uncommitted (M0) to anti-inflammatory (M2) phenotypes. To test this hypothesis, we quantified cytokine secretion in macrophages under different matrix conditions after 72 hours. **Figure 2E** shows distinct cytokine profiles: macrophages on random matrices secreted relatively high levels of proinflammatory cytokines, such as MCP-1, IFN-γ, IL-12p70, IL-1β, and TNF-α. Conversely, macrophages on crosslinked and aligned matrices demonstrated reduced secretion of these cytokines. Moreover, anti-inflammatory cytokines such as IL-4, IL-10, and TGF-β1 were more prevalent in macrophages on both stiff matrices, with slightly greater secretion in crosslinked matrices. These cytokine profiles suggest that macrophages on both crosslinked and aligned matrices exhibit a reduced inflammatory response, as evidenced by decreased secretion of proinflammatory cytokines and increased secretion of anti-inflammatory cytokines. These results align cytokine secretion profiling with immunophenotyping **(Figure 2C and 2E)**.

In conclusion, our findings demonstrate that matrix stiffening through crosslinking and alignment promotes the polarization of uncommitted macrophages (M0) toward an anti-inflammatory M2 phenotype, specifically M2a, characterized by increased expression of CD105 and CD206 [32]. This shift in the macrophage phenotype, accompanied by reduced secretion of proinflammatory cytokines and increased secretion of anti-inflammatory cytokines, underscores the influence of the physical properties of the TME on immune cell behavior.

### 3.3. RNA sequencing reveals similar transcriptomes and increased PI3K signaling in macrophages cultured on stiff matrices

To determine how the matrix influences the polarization of macrophages toward an anti-inflammatory phenotype, we conducted a functional transcriptome analysis using RNA sequencing. As depicted in **Figure 3A**, our findings indicate that the gene expression patterns of macrophages cultured on stiff matrices, whether achieved through crosslinking or collagen fibril alignment, were similar. In contrast, macrophages cultured on a relatively more compliant random matrix presented a distinct transcriptome pattern. These findings suggest that matrix stiffness plays a crucial role in shaping the gene expression landscape of macrophages. We further substantiated these results through principal component analysis (PCA) and a correlation matrix, as presented in **Figure 3B and 3C**, respectively. The PCA plot illustrates a clear separation between the random, crosslinked, and aligned conditions, with the first two principal components explaining a significant portion of the variance in the data. The correlation matrix supported this finding, showing high correlations within the stiff matrix conditions and lower correlations with the random condition, indicating distinct gene expression profiles.

**Figure 3:**
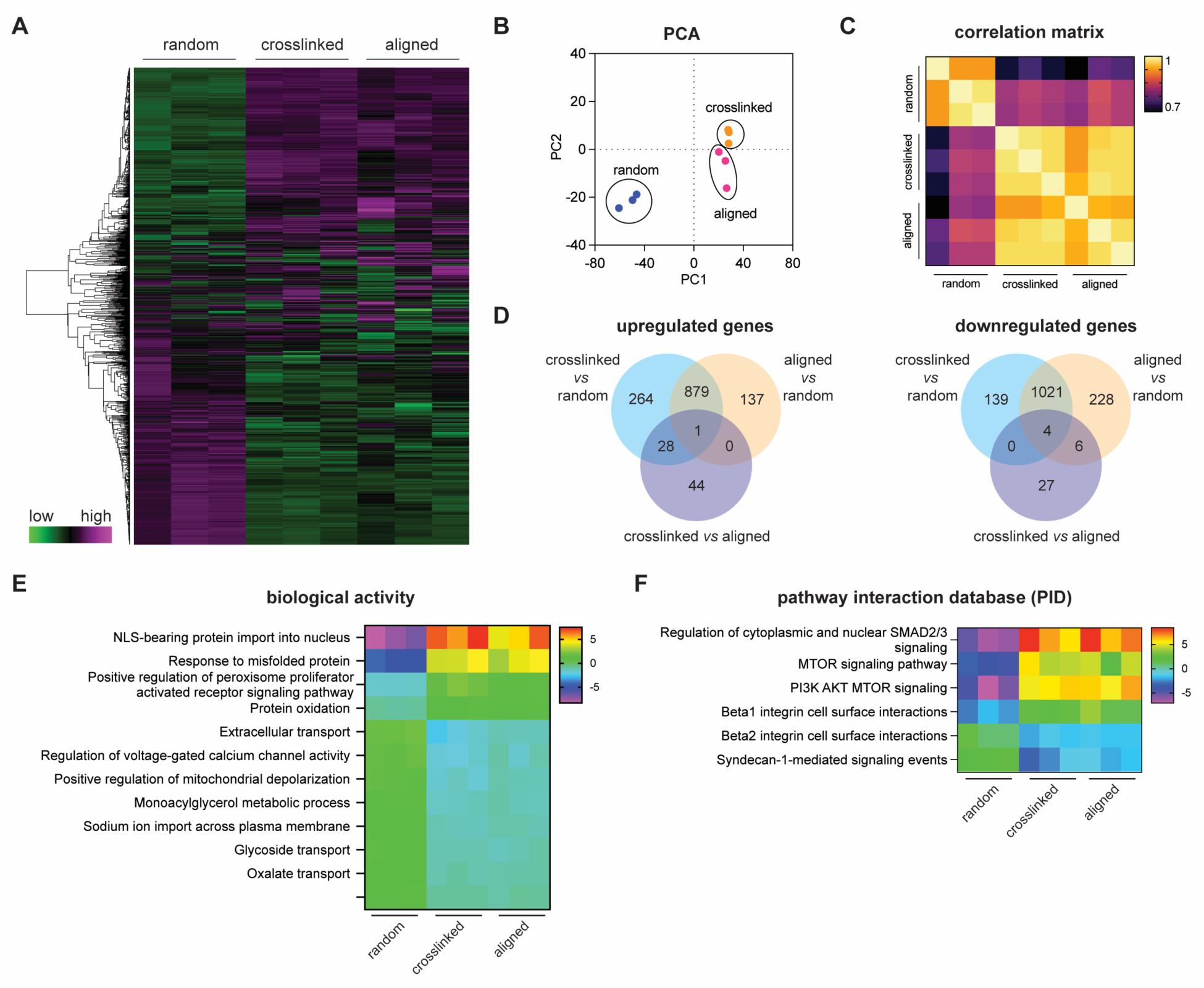
Transcriptome analysis of macrophages cultivated in different matrices. Functional transcriptome analysis was conducted using the iDEP web-based tool. **(A)** Heatmap displaying the overall gene expression levels in macrophages in various matrices (green represents low expression; violet represents high expression). **(B)** Principal component analysis (PCA) showing the separation of gene expression profiles between the different conditions. **(C)** Correlation matrix of gene expression data illustrating the relationships between samples. **(D)** Venn diagrams illustrating the up- and downregulated DEGs with an FDR cutoff of 0.05 and a minimum fold change of 2. **(E)** Heatmap of enriched biological activities and **(F)** curated pathway interaction database (PID) pathways, with blue representing low enrichment and red representing high enrichment. Three independent experiments were used for RNA sequencing.

From the RNA sequencing data, we proceeded to analyze differentially expressed genes (DEGs) using a false discovery rate (FDR) threshold of 0.05 and a minimum fold change of 2 for all conditions compared with cells cultured on a random compliant matrix. As illustrated in **Figure 3D**, we visualized the up- and downregulated DEGs via a Venn diagram. Compared with the random conditions, both the crosslinked and aligned conditions presented a significant number of DEGs, indicating substantial transcriptional reprogramming in response to matrix stiffness. The overlap between crosslinked and aligned conditions suggests that these changes are predominantly driven by matrix stiffness rather than specific structural features.

To predict enrichment in biological activities and pathway interactions, we conducted parametric gene set enrichment analysis (PGSEA). This analysis allowed us to predict biological pathways without bias from the identified DEGs, as DEGs heavily depend on the specific criteria set for the cutoff, in our case, an FDR cutoff of 0.05 and a minimum fold change of 2. As illustrated in **Figure 3E**, we observed greater gene set enrichment in biological processes such as NLS-bearing protein import into the nucleus, response to misfolded proteins, positive regulation of the peroxisome proliferator-activated receptor signaling pathway, and protein oxidation in both stiff matrices (crosslinked and aligned conditions). These processes collectively enhance the ability of M2 macrophages to resolve inflammation, facilitate tissue remodeling, and sustain anti-inflammatory activities. Previous reports have indicated that positive regulation of the peroxisome proliferator-activated receptor signaling pathway promotes macrophage polarization toward anti-inflammatory phenotypes [53–55]. The enrichment of this pathway in stiff matrices supports our hypothesis that matrix stiffness promotes an anti-inflammatory macrophage phenotype.

According to the analysis of curated pathway interaction databases **(Figure 3F)**, we predict that the regulation of cytoplasmic and nuclear SMAD2/3, the MTOR signaling pathway, the PI3K-AKT-MTOR signaling pathway, and the beta1 integrin cell surface interaction pathway are upregulated in stiff matrices (both crosslinked and aligned samples). These pathways are known to play crucial roles in cellular signaling and mechanotransduction. Based on these functional transcriptome analyses, we hypothesize that the activation of the JAK-STAT signaling pathway may be due to the increased secretion of IL-4, as illustrated in **Figure 2E**, which in turn polarizes M0 macrophages toward the M2a phenotype. On the other hand, beta1 integrin and PI3K-AKT-MTOR signaling might contribute to the activation of mechanotransduction pathways, leading to IL-4 secretion. Previous studies have shown that beta1 integrin and PI3K-AKT-MTOR activation is a crucial step in the M2 activation of macrophages in response to IL-4 [56–58].

### 3.4. Inhibiting the integrin β1 and PI3K pathways reduces M2 polarization in stiff matrices

To validate the hypotheses arising from the predicted pathway interactions from RNA sequencing analysis, we inhibited possible signaling pathways involved in mechanosensing and transduction using an antibody against integrin β1 (anti-integrin β1) and chemical inhibitors against PI3K (LY294002), ROCK (Y-27632), YAP (verteporfin), and Piezo1 (GsMTx4) signaling. M0 macrophages were cultured in the presence or absence of the antibody or chemical inhibitors for 3 days. As shown in **Figure 4A**, we found a significant decrease in the percentage of CD206-positive cells when integrin β1 and the PI3K pathway were blocked, whereas other inhibitors did not significantly change the percentage of CD206-positive cells. This finding led to the hypothesis that integrin β1 and PI3K signaling are associated with M2a polarization in stiffened matrices. Our findings are in line with other studies showing the involvement of integrin β1 in M2 polarization [59].

**Figure 4:**
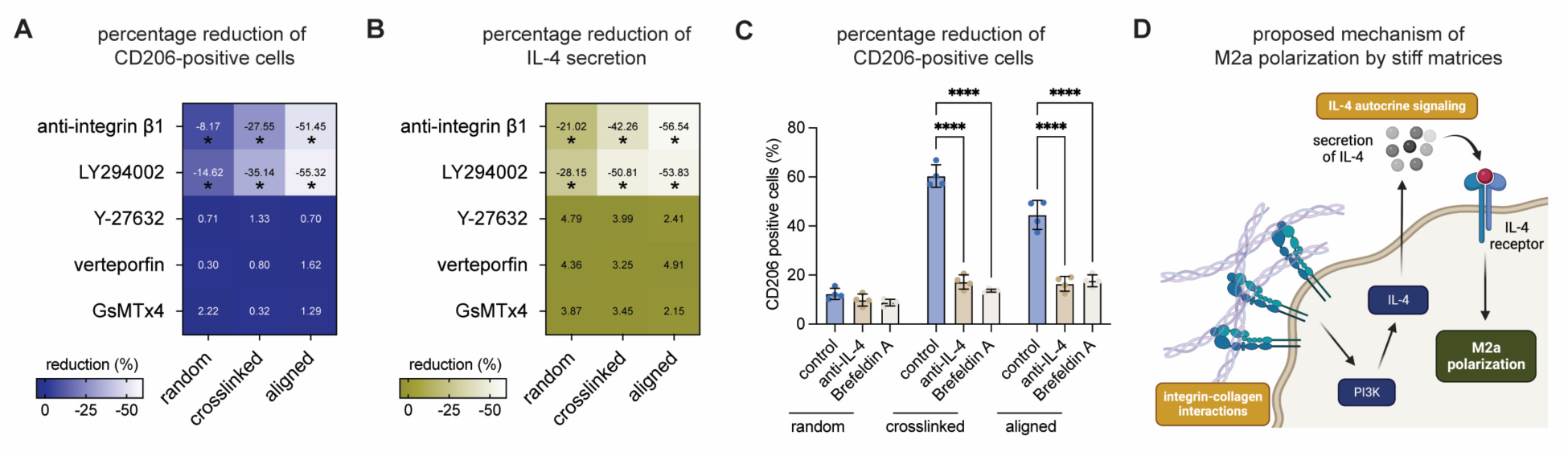
Inhibition of mechanotransduction pathways and proposed mechanism of matrix stiffness-mediated M2 polarization. **(A)** Quantitative analysis of the percentage reduction in CD206-positive cells after treatment with different inhibitors (anti-integrin β1, LY294002, Y-27632, verteporfin, and GsMTx4) across various matrix conditions. **(B)** Quantitative analysis of the percentage reduction in IL-4 secretion in cell culture media after treatment with different inhibitors. The data were normalized to those of cells cultured on each matrix without treatment, and a significance test was performed against the untreated control. **(C)** Quantitative analysis of CD206-positive cells after neutralizing IL-4 with an antibody or blocking cytokine secretion with brefeldin A. **(D)** Proposed mechanism of M2a polarization by stiffened matrices. Levels of statistical significance were assessed using two-way ANOVA followed by Tukey’s post hoc test. Significance levels were established as follows: *p < 0.05, **p < 0.01, ***p < 0.001, ****p < 0.0001. Four independent replicates of the experiments were performed.

As the differentiation of M2a macrophages requires IL-4 or IL-4 in combination with IL-13 [32, 39, 59, 60], we further analyzed IL-4 in the cell culture supernatant and found a significant reduction in IL- 4 secretion when integrin β1 or the PI3K pathway was blocked (**Figure 4B**), further confirming that both pathways are involved in regulating M2a polarization. To demonstrate that secreted IL-4 might act in an autocrine fashion to polarize M0 toward the M2a subtype, we neutralized secreted IL-4 by supplementing the cell culture medium with an anti-IL-4 antibody or brefeldin A to inhibit the secretion of cytokines. As a result of these treatments, there was a significant reduction in the number of CD206-positive cells (**Figure 4C**), suggesting that IL-4 is secreted by cells and acts in an autocrine fashion to modulate M2a polarization.

Based on our data, we propose that mechanomodulation of M2a polarization occurs through a stiffened matrix via integrin β1 and PI3K signaling. Upon mechanosensing, cells increase the production and secretion of IL-4, which subsequently acts in an autocrine manner during macrophage polarization toward the M2a subtype. This proposed mechanism is depicted in **Figure 4D**.

### 3.5. Crosslinked and aligned matrices reduced the proinflammatory response in macrophages

The inflammatory mediators within the tumor microenvironment are known to drive malignancy and modulate cell polarization [3, 61, 62]. Interestingly, anti-inflammatory macrophages are predominant in these microenvironments, presenting a paradox in the immune landscape [63]. We thus investigated whether inducing a proinflammatory response in macrophages cultured on stiffened matrices could effectively shift their phenotype to a more proinflammatory state (M1).

To investigate this possibility, we induced macrophage polarization toward proinflammatory phenotypes using IFN-γ and LPS for 48 hours (**Figure 5A)**. Macrophage polarization was assessed by quantifying the expression of HLADR-positive cells, a well-established marker of M1 macrophages [25, 39], and by measuring the secretion of proinflammatory cytokines. The results demonstrated that macrophages cultured on crosslinked and aligned matrices presented fewer HLADR-positive cells than those cultured on a random matrix **(Figure 5B)**. The reduced secretion of IL-1β, MCP-1, and TNF-α in macrophages cultured on crosslinked and aligned matrices suggested that the mechanical stiffness of the ECM may inhibit the proinflammatory functions of these cells **(Figure 5C)**. These cytokines are crucial for initiating and sustaining inflammatory responses, which are essential for antitumor immunity. The unchanged levels of IFN-γ and IL-12p70 across different substrate conditions might indicate that certain cytokine pathways are less sensitive to mechanical cues or that compensatory mechanisms maintain their expression. The suppression of macrophage inflammatory responses in stiffened environments could contribute to a more immunosuppressive tumor microenvironment, which favors tumor growth and progression.

**Figure 5:**
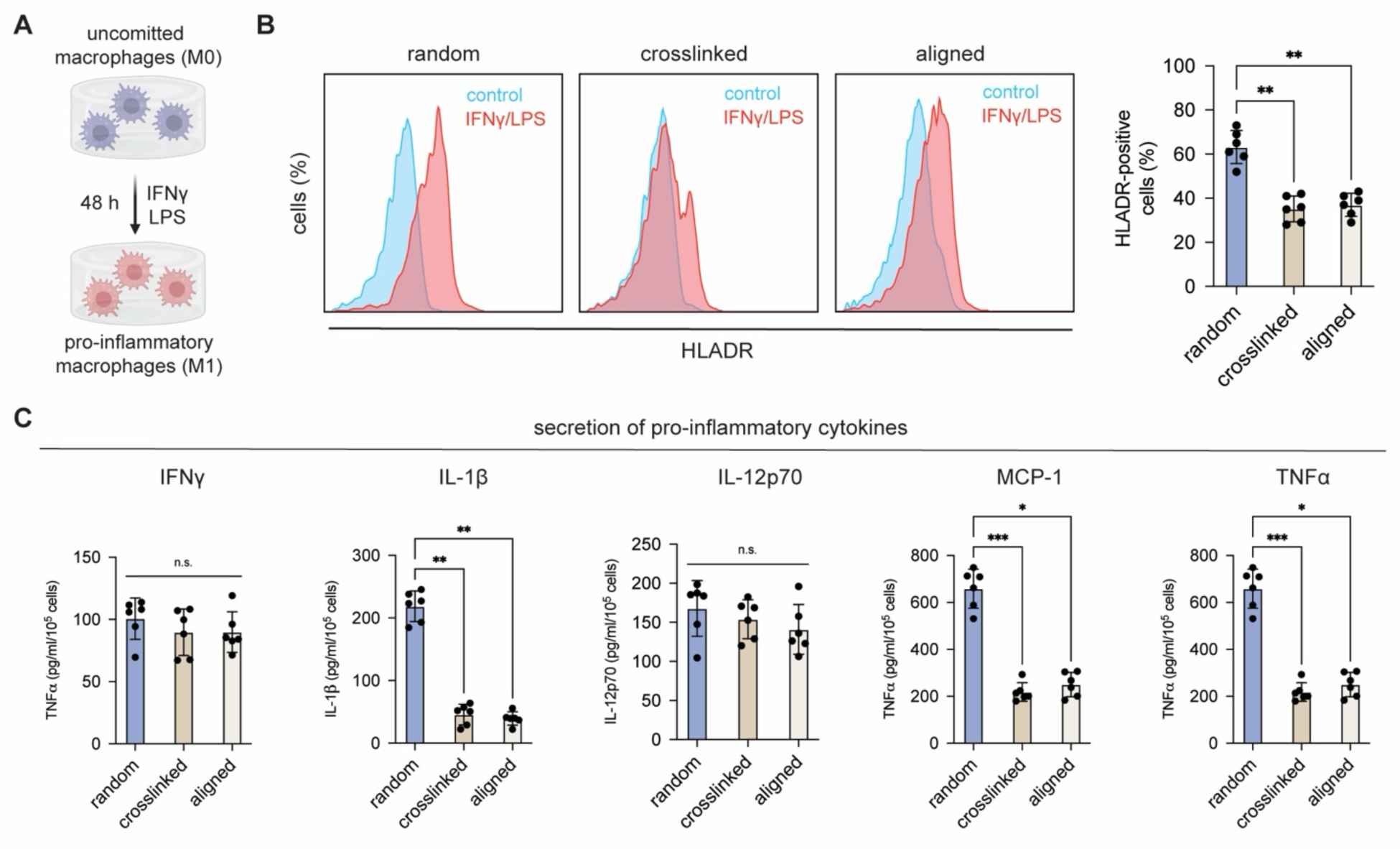
Induction of proinflammatory macrophage phenotypes in different matrices. **(A)** Schematic illustration of macrophage polarization toward a proinflammatory phenotype after treatment with IFN-γ and LPS for 48 hours. **(B)** Representative histogram plots of HLA-DR expression in macrophages treated with or without IFN-γ and LPS, accompanied by quantitative analysis of HLA-DR-positive cells under various matrix conditions. **(C)** Secretion of proinflammatory cytokines, namely, IL-1β, IL-12p70, MCP-1, and TNFα, was quantified using a custom-made multiplex bead-based ELISA. The data are presented as the means ± SDs. Levels of statistical significance were assessed using one-way ANOVA followed by Tukey’s post hoc test. Significance levels were established as follows: *p < 0.05, **p < 0.01, ***p < 0.001, ****p < 0.0001. The experiments were performed in 6 independent replicates.

Overall, the observation that stiffened matrices suppress macrophage inflammatory responses highlights the complex interplay between mechanical signals and immune cell function within the TME. Additionally, our results highlight the need for therapeutic strategies that can target the ECM to restore or enhance macrophage proinflammatory activity. Understanding the molecular mechanisms by which matrix stiffness influences macrophage function could lead to the development of therapies aimed at modulating the physical properties of the TME.

### 3.6. Matrix alignment and macrophage coculture significantly impact cancer cell behavior

To better understand the complex interactions among cancer cells, macrophages, and their microenvironment, we conducted coculture experiments to observe their influence on cancer cell behavior. Specifically, we investigated how different matrix conditions affect two breast cancer cell lines, MDA-MB-231 and MCF-7, in both monoculture and coculture with M0 macrophages, with a focus on their proliferation, infiltration, and cytokine secretion profiles.

**Figure 6A** shows images of CSFE-stained MDA-MB-231 cells (green) and Hoechst 33342-stained macrophage nuclei (blue) in various matrix conditions. These images reveal that matrix alignment influences the cell distribution and morphology. In monoculture, cancer cells are evenly spread in random and crosslinked matrices but align along the fibers in the aligned matrix. In coculture, macrophages interact closely with cancer cells but do not affect their distribution. By quantifying the number of MDA-MB-231 cells **(Figure 6B)**, we found that in monocultures, the number of cells was significantly greater in crosslinked and aligned matrices than in random matrices, suggesting increased proliferation. Under coculture conditions, the number of cancer cells was significantly greater than that under monoculture conditions, especially in aligned matrices, indicating that macrophages may promote cancer cell proliferation. We also measured the percentage of infiltrating cancer cells **(Figure 6C)**. The results revealed higher infiltration rates in coculture conditions across all matrix conditions, with the highest rates in aligned matrices, highlighting the impact of matrix alignment on cancer cell invasion. Cytokine secretion profiles **(Figure 6D)**, analyzed using bead-based multiplex ELISA, revealed that the levels of the anti-inflammatory cytokines IL-4 and IL-10 were significantly elevated in coculture conditions featuring aligned and crosslinked matrices. Conversely, the levels of the proinflammatory cytokines IL-1β and TNF-α are markedly greater in coculture settings with aligned matrices. Additionally, cytokines related to immune activation and recruitment, such as MCP-1, IL-2, IL-8, and IP-10, exhibit increased secretion under coculture conditions with aligned and crosslinked matrices. Other cytokines, including IL-6, IL-12p70, IFN-γ, IL-17A, and TGF-β1, exhibit variable levels across different conditions. Further investigation of macrophage phenotypes revealed a greater percentage of HLA-DR-positive macrophages in coculture conditions, particularly in aligned matrices, indicating enhanced macrophage activation **(Figure 6E)**. Additionally, there were greater numbers of CD206-positive macrophages in coculture conditions, suggesting an increase in macrophage polarization toward the M2 phenotype **(Figure 6B)**.

**Figure 6:**
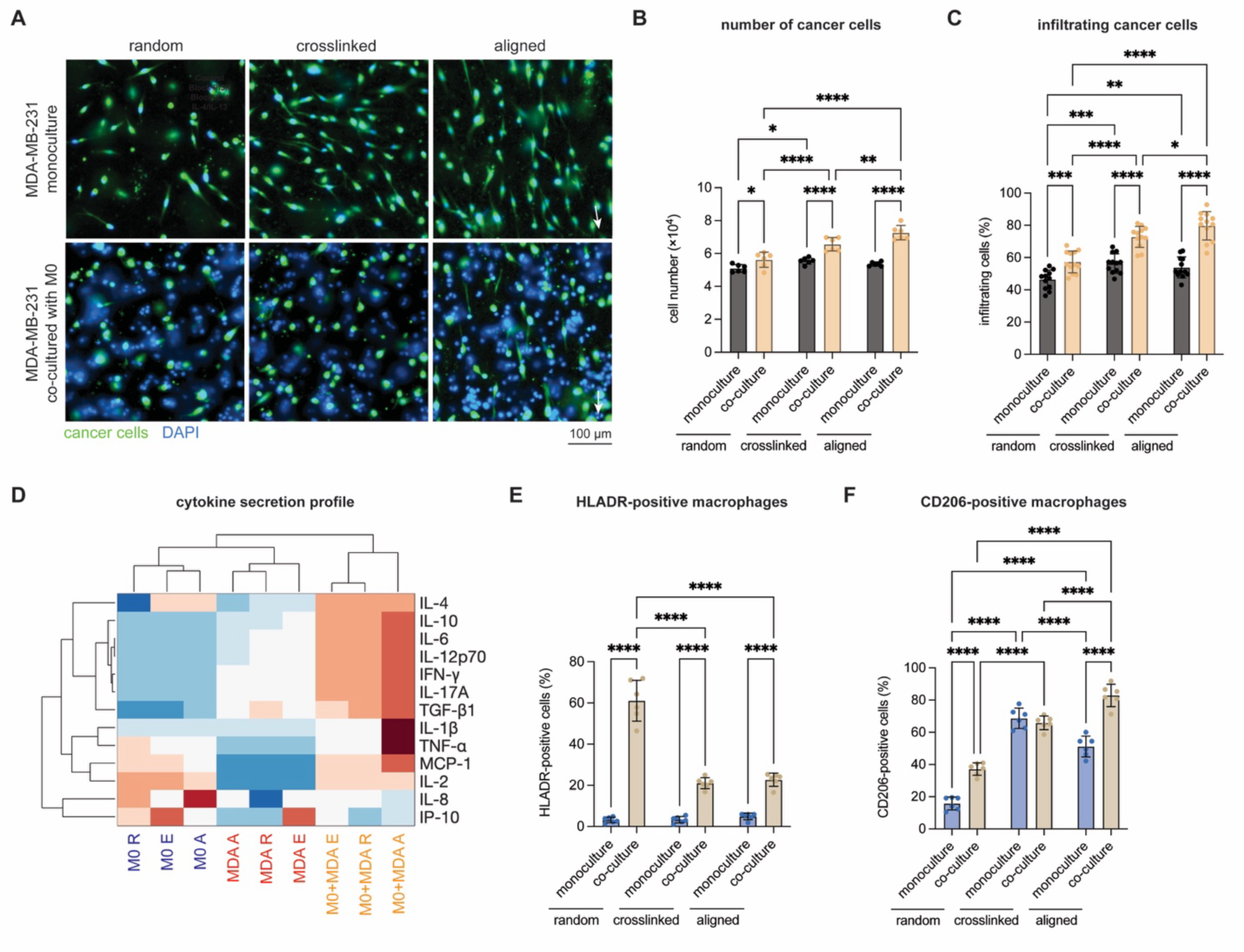
Macrophage and MDA-MB-321 breast cancer cell interactions in different matrices. **(A)** Representative fluorescence images showing CSFE-stained cancer cells (green) and Hoechst 33342-stained nuclei of macrophages (blue) under different matrix conditions in monoculture and coculture. White arrows indicate the direction of fibril alignment. (Scale bar = 100 µm) **(B)** Quantification of the number of cancer cells. **(C)** Percentages of infiltrating cancer cells under different matrix conditions in monocultures and cocultures. **(D)** Heatmap showing cytokine secretion profiles of M0 macrophages and MDA-MB-231 cells under various matrix conditions, including random (r), crosslinked (r), and aligned (a) matrices in monoculture and coculture. **(E)** Percentages of HLA-DR-positive macrophages in monoculture and coculture conditions under different matrix conditions. **(F)** Percentage of CD206-positive macrophages in monoculture and coculture conditions under different matrix conditions. The data are presented as the means ± SDs. Levels of statistical significance were assessed using two-way ANOVA followed by Tukey’s post hoc test. Significance levels were established as follows: *p < 0.05, **p < 0.01, ***p < 0.001, ****p < 0.0001. The experiments were performed in 6 independent replicates.

We next investigated the effects of different matrices on MCF-7 cancer cells and their interactions with M0 macrophages. **Figure 7A** shows images of CFSE-stained MCF-7 cells (green) and Hoechst 33342- stained macrophage nuclei (blue) in various matrix conditions. Quantitative analysis of cell numbers revealed that compared with the random matrix, both the crosslinked and aligned matrices increased the number of cancer cells under both monoculture and coculture conditions **(Figure 7B)**. Additionally, the percentage of infiltrating cancer cells was significantly greater under coculture conditions, especially in the aligned matrices **(Figure 7C)**. The cytokine secretion profile of macrophages cocultured with MCF-7 cancer cells varied significantly depending on the matrix conditions, as illustrated in the heatmap **(Figure 7D)**. The aligned matrices induced a more pronounced proinflammatory cytokine response, characterized by elevated levels of TNF-α, IL-1β, and IL-12p70. These matrices also show increased levels of IL-4 and IL-2. In contrast, crosslinked and random matrices promote a more regulated or anti-inflammatory cytokine profile, marked by increased expression of IL-10 and IL-6. Further analysis of macrophage polarization revealed that the percentage of HLA-DR-positive macrophages across all matrix types was significantly greater under coculture conditions than under monoculture conditions, indicating that the presence of MCF- 7 cells influences the polarization of M1 macrophages **(Figure 7E)**. Similarly, the percentage of CD206- positive macrophages was greater under coculture conditions than under monoculture conditions, particularly in the aligned matrices **(Figure 7F)**.

**Figure 7:**
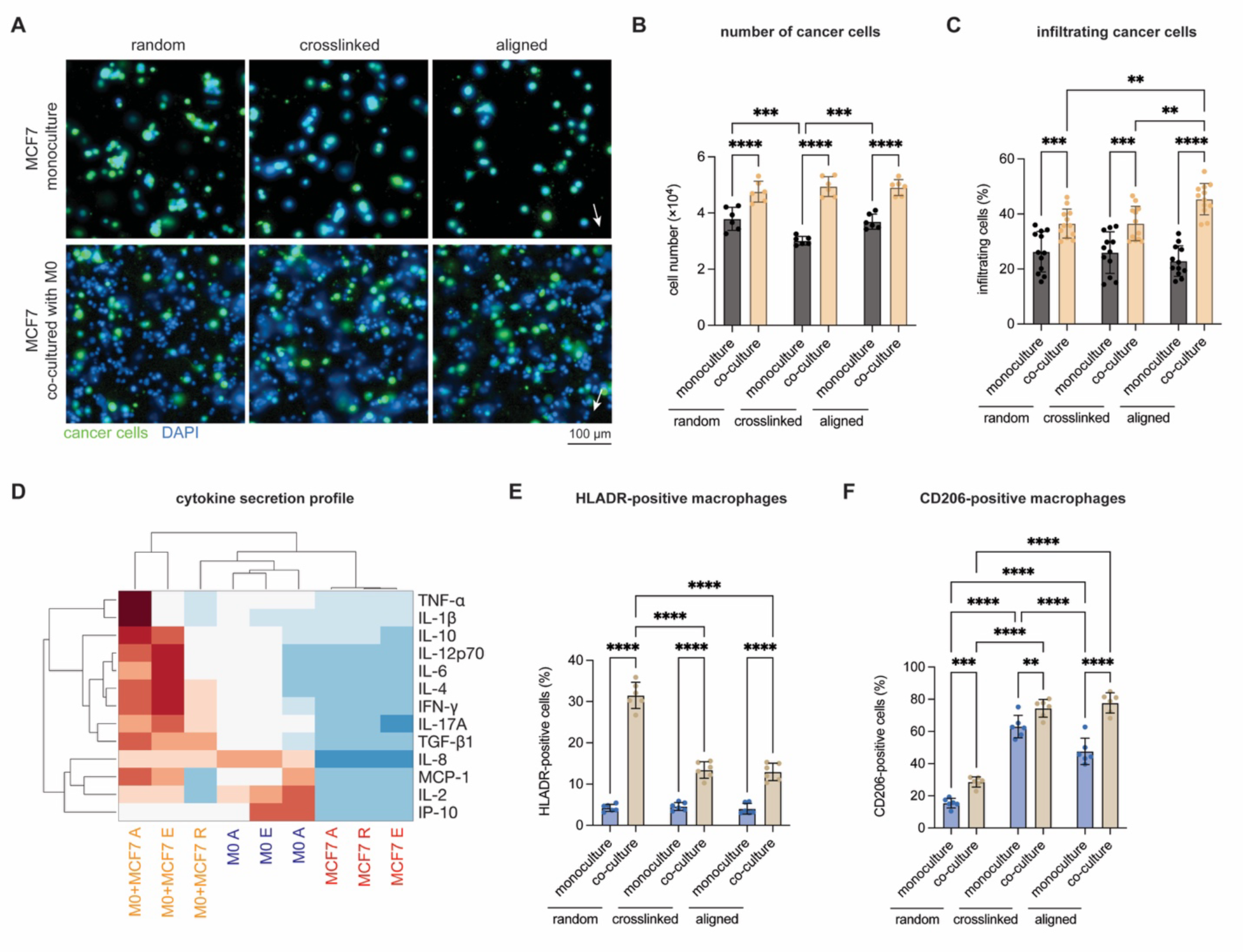
Macrophage and MCF-7 breast cancer cell interactions in different matrices. **(A)** Representative fluorescence images showing CSFE-stained cancer cells (green) and Hoechst 33342-stained nuclei of macrophages (blue) in different matrix conditions in monoculture and coculture. White arrows indicate the direction of fibril alignment. (Scale bar = 100 µm) **(B)** Quantification of the number of cancer cells. **(C)** Percentages of infiltrating cancer cells under different matrix conditions in monocultures and cocultures. **(D)** Heatmap showing cytokine secretion profiles of M0 macrophages and MCF-7 cells under various matrix conditions, including random (r), crosslinked (r), and aligned (a) matrices in monoculture and coculture. **(E)** Percentages of HLA-DR-positive macrophages in monoculture and coculture conditions under different matrix conditions. **(F)** Percentage of CD206-positive macrophages in monoculture and coculture conditions under different matrix conditions. The data are presented as the means ± SDs. Levels of statistical significance were assessed using two-way ANOVA followed by Tukey’s post hoc test. Significance levels were established as follows: *p < 0.05, **p < 0.01, ***p < 0.001, ****p < 0.0001. The experiments were performed in 6 independent replicates.

Comparing MCF-7 and MDA-MB-231 cells, coculture with macrophages significantly increased the number of cancer cells, particularly in aligned matrices, indicating that macrophages may promote cancer cell proliferation. In both cell types, coculture, especially with aligned and crosslinked matrices, elevated levels of anti-inflammatory cytokines (e.g., IL-4 and IL-10) and proinflammatory cytokines (e.g., IL-1β and TNF-α) suggest that specific matrix alignments create complex immune environments. Higher percentages of HLA-DR-positive and CD206-positive macrophages were observed in coculture with aligned matrices for both MCF-7 and MDA-MB-231 cells, indicating a shift toward an M2 phenotype and the promotion of an anti-inflammatory milieu. The increased levels of proinflammatory cytokines and HLA-DR-positive cells in coculture with aligned matrices, despite an overall tumor-supportive environment, highlight the dual roles of proinflammatory cytokines such as IL-1β and TNF-α [62]. These cytokines can promote tumor growth and survival while also stimulating immune responses to target cancer cells [64, 65]. The presence of HLA-DR-positive macrophages indicates an activated state that enhances the recruitment and activation of other immune cells, potentially fostering an antitumor response [23]. However, tumors can exploit this immune activation to create a chronic inflammatory state that supports tumor growth and metastasis [62]. The proinflammatory environment created by macrophages and aligned matrices may contribute to an immune landscape that, while potentially targeting the tumor, ultimately supports tumor progression through chronic inflammation and immune evasion mechanisms [35, 66, 67]. This paradox suggests that the TME is finely balanced, with both pro- and anti-inflammatory signals playing critical roles in cancer progression.

## 4. Conclusions

This study provides valuable insights into how the mechanical properties of the ECM within the TME influence macrophage behavior and cancer progression. Using 3D collagen matrices with random, crosslinked, and aligned fibrils, we mimicked the mechanical environment of cancer tissues, revealing significant biological responses. Increased ECM stiffness, achieved through chemical crosslinking or collagen fibril alignment, promotes macrophage polarization toward the tumor-supportive M2 phenotype. M2 macrophages, characterized by increased CD105 and CD206 expression and reduced proinflammatory cytokine secretion, contribute to a tumor-promoting environment. We identified the integrin β1 and PI3K signaling pathways as critical mediators of this process. Blocking these pathways reduces M2 polarization and IL-4 secretion, indicating their crucial roles in mechanotransduction and autocrine signaling. Transcriptome analysis further supported these findings. Compared with those cultured on compliant matrices, macrophages cultured on stiff matrices presented distinct gene expression profiles, with enriched biological processes related to immune suppression and tissue remodeling. The upregulated pathways, such as the PI3K-AKT-MTOR and integrin signaling pathways, suggest that these pathways are vital for macrophage mechanosensing and phenotypic modulation. In coculture experiments with breast cancer cell lines (MDA-MB-231 and MCF-7), we observed that macrophages within stiffened matrices enhanced cancer cell proliferation and invasion. Elevated levels of both anti-inflammatory and proinflammatory cytokines in these conditions indicate a complex immune environment that facilitates tumor progression and immune evasion, underscoring the importance of ECM stiffness in promoting a tumor-supportive microenvironment. These results highlight several potential therapeutic strategies. Modifying ECM stiffness to disrupt the supportive environment for M2 macrophages could inhibit tumor progression [68]. Reprogramming macrophages from an M2 phenotype to an M1 phenotype could enhance antitumor immunity [52]. Inhibitors targeting the integrin β1 and PI3K pathways might prevent macrophage polarization toward a tumor-supportive phenotype. Overall, understanding and targeting the interactions between ECM stiffness and macrophage polarization are promising approaches for cancer therapies.

## Acknowledgments

The authors acknowledge support from the New York University Abu Dhabi (NYUAD) Faculty Research Fund (AD266). The authors would also like to acknowledge support from the NYUAD core technology platform: Light Microscopy, Molecular and Cell Biology, and Sequencing.

## Author statement

The authors confirm that neither the manuscript nor its contents have been previously published in any other form and are not intended for publication in any other journal at the time of submission. All the authors have read and approved the submission of the revised manuscript.

## CRediT authorship contribution statement

**Jiranuwat Sapudom:** Writing – review & editing, Writing – original draft, Investigation, Formal analysis, Conceptualization. **Paul Tipay:** Writing – review & editing, Investigation, Formal analysis. **Jeremy Teo:** Writing – review & editing, Investigation, Formal analysis, Conceptualization, Supervision and Funding acquisition.

## Data availability

All the data and custom MATLAB scripts used in this study are available upon request from the authors. The MATLAB script for network topology analysis is freely accessible at https://git.sc.uni-leipzig.de/pe695hoje/topology-analysis.

## Supplementary methods

### Processing of RNA Sequencing data

The quality of the raw FASTQ-sequenced reads was initially assessed using FastQC v0.11.5 (available online at http://www.bioinformatics.babraham.ac.uk/projects/fastqc/). Next, the reads were processed through Trimmomatic v0.36 (10.1093/bioinformatics/btu170) for quality trimming and adapter sequence removal using the following parameters: ILLUMINACLIP.fa:2:30:10, TRAILING:3, LEADING:3, SLIDINGWINDOW:4:15, and MINLEN:36. The trimmed read pairs were then processed with Fastp (10.1093/bioinformatics/bty560) to remove poly-G tails and Novaseq/NextSeq-specific artifacts. After quality trimming, the reads were reassessed using FastQC. Subsequently, the reads were aligned to the human reference genome GRCh38.p4 using HISAT2 (10.1038/nmeth.3317) with the default parameters and the –dta flag. The resulting SAM alignments were converted to BAM format and sorted by coordinates using SAMtools v1.3.1 (10.1093/bioinformatics/btp352). The sorted alignment files were then processed with HTSeq-count v0.6.1p1 (10.1093/bioinformatics/btu638) using the options -s no -t exon -I gene_id to generate raw counts. Concurrently, the sorted alignments were processed through Stringtie v1.3.0 (10.1038/nprot.2016.095) for transcriptome quantification. The process included running Stringtie, then Stringtie merge to create a merged transcriptome GTF file for all samples, followed by another round of Stringtie using the merged GTF file. Finally, RNA-Seq-specific QC metrics per sample were generated using Qualimap v2.2.2 (10.1093/bioinformatics/bts503). The RNA-Seq data were merged using the NASQAR toolbox, which is publicly accessible at http://nasqar.abudhabi.nyu.edu/ (10.1186/s12859-020-03577-4).

---

**Table S1:** Antibodies used in the study. All antibodies were purchased from Biolegend.

| Marker | Color/Format | Host/Target | Isotype | Clone | Catalog Nr. |
| --- | --- | --- | --- | --- | --- |
| CD105 | Alexa Fluor 647 | Mouse anti-Human | IgG1 κ | 43A3 | 323212 |
| CD163 | PE-Cy7 | Mouse anti-Human | IgG1 κ | GHI/61 | 333614 |
| CD206 | Brilliant Violet 510 | Mouse anti-Human | IgG1 κ | 15-2 | 321138 |
| HLA-DR | Alexa Fluor 488 | Mouse anti-Human | IgG2a κ | L243 | 307656 |

## References

[1] X. Zhang, H. Ma, Y. Gao, Y. Liang, Y. Du, S. Hao, T. Ni, The Tumor Microenvironment: Signal Transduction, Biomolecules 14(4) (2024) 10.3390/biom14040438.

[2] A. Mancini, M.T. Gentile, F. Pentimalli, S. Cortellino, M. Grieco, A. Giordano, Multiple aspects of matrix stiffness in cancer progression, Front Oncol 14 (2024) 1406644, 10.3389/fonc.2024.1406644.

[3] T.R. Mempel, J.K. Lill, L.M. Altenburger, How chemokines organize the tumour microenvironment, Nat Rev Cancer 24(1) (2024) 28–50, 10.1038/s41568-023-00635-w.

[4] A. Akinpelu, T. Akinsipe, L.A. Avila, R.D. Arnold, P. Mistriotis, The impact of tumor microenvironment: unraveling the role of physical cues in breast cancer progression, Cancer Metastasis Rev 43(2) (2024) 823–844, 10.1007/s10555-024-10166-x.

[5] S. Benmelech, T. Le, M. McKay, J. Nam, K. Subramaniam, D. Tellez, G. Vlasak, M. Mak, Biophysical and biochemical aspects of immune cell-tumor microenvironment interactions, APL Bioeng 8(2) (2024) 021502, 10.1063/5.0195244.

[6] Y. Xiao, Z. Wang, M. Gu, P. Wei, X. Wang, W. Li, Cancer-associated fibroblasts: heterogeneity and their role in the tumor immune response, Clin Exp Med 24(1) (2024) 126, 10.1007/s10238-024-01375-3.

[7] R. Kalluri, The biology and function of fibroblasts in cancer, Nat Rev Cancer 16(9) (2016) 582–598, 10.1038/nrc.2016.73.

[8] C.P. El-Haibi, G.W. Bell, J. Zhang, A.Y. Collmann, D. Wood, C.M. Scherber, E. Csizmadia, O. Mariani, C. Zhu, A. Campagne, M. Toner, S.N. Bhatia, D. Irimia, A. Vincent-Salomon, A.E. Karnoub, Critical role for lysyl oxidase in mesenchymal stem cell-driven breast cancer malignancy, Proc Natl Acad Sci U S A 109(43) (2012) 17460–17465, 10.1073/pnas.1206653109.

[9] W. Han, S. Chen, W. Yuan, Q. Fan, J. Tian, X. Wang, L. Chen, X. Zhang, W. Wei, R. Liu, J. Qu, Y. Jiao, R.H. Austin, L. Liu, Oriented collagen fibers direct tumor cell intravasation, Proc Natl Acad Sci U S A 113(40) (2016) 11208–11213, 10.1073/pnas.1610347113.

[10] M.W. Conklin, J.C. Eickhoff, K.M. Riching, C.A. Pehlke, K.W. Eliceiri, P.P. Provenzano, A. Friedl, P.J. Keely, Aligned collagen is a prognostic signature for survival in human breast carcinoma, Am J Pathol 178(3) (2011) 1221–1232, 10.1016/j.ajpath.2010.11.076.

[11] B.C. Quartey, G. Torres, M. ElGindi, A. Alatoom, J. Sapudom, J.C.M. Teo, Tug of war: Understanding the dynamic interplay of tumor biomechanical environment on dendritic cell function, Mechanobiology in Medicine 2(3) (2024) 100068, 10.1016/j.mbm.2024.100068.

[12] A. Alatoom, M. ElGindi, J. Sapudom, J.C.M. Teo, The T Cell Journey: A Tour de Force, Adv Biol (Weinh) 7(1) (2023) e2200173, 10.1002/adbi.202200173.

[13] M. Golo, P.L.H. Newman, D. Kempe, M. Biro, Mechanoimmunology in the solid tumor microenvironment, Biochem Soc Trans 52(3) (2024) 1489–1502, 10.1042/BST20231427.

[14] P. Riedl, J. Sapudom, C. Clemens, L. Orgus, A. Proger, J.C.M. Teo, T. Pompe, Phenotype Switching of Breast Cancer Cells upon Matrix Interface Crossing, ACS Appl Mater Interfaces 15(20) (2023) 24059–24070, 10.1021/acsami.3c01099.

[15] J. Sapudom, S. Karaman, B.C. Quartey, W.K.E. Mohamed, N. Mahtani, A. Garcia-Sabate, J. Teo, Collagen Fibril Orientation Instructs Fibroblast Differentiation Via Cell Contractility, Adv Sci (Weinh) 10(22) (2023) e2301353, 10.1002/advs.202301353.

[16] B.R. Seo, X. Chen, L. Ling, Y.H. Song, A.A. Shimpi, S. Choi, J. Gonzalez, J. Sapudom, K. Wang, R.C. Andresen Eguiluz, D. Gourdon, V.B. Shenoy, C. Fischbach, Collagen microarchitecture mechanically controls myofibroblast differentiation, Proc Natl Acad Sci U S A 117(21) (2020) 11387–11398, 10.1073/pnas.1919394117.

[17] J. Sapudom, A. Alatoom, P. Tipay, J.C.M. Teo, Matrix alignment and density modulate YAP- mediated T-cell immune suppression, bioRxiv (2024) 2024.2003.2019.585707, 10.1101/2024.03.19.585707.

[18] P.P. Provenzano, K.W. Eliceiri, J.M. Campbell, D.R. Inman, J.G. White, P.J. Keely, Collagen reorganization at the tumor-stromal interface facilitates local invasion, BMC Med 4(1) (2006) 38, 10.1186/1741-7015-4-38.

[19] K.M. Riching, B.L. Cox, M.R. Salick, C. Pehlke, A.S. Riching, S.M. Ponik, B.R. Bass, W.C. Crone, Y. Jiang, A.M. Weaver, K.W. Eliceiri, P.J. Keely, 3D collagen alignment limits protrusions to enhance breast cancer cell persistence, Biophys J 107(11) (2014) 2546–2558, 10.1016/j.bpj.2014.10.035.

[20] I.M. Joshi, M. Mansouri, A. Ahmed, D. De Silva, R.A. Simon, P. Esmaili, D.E. Desa, T.M. Elias, E.B. Brown, 3rd, V.V. Abhyankar, Microengineering 3D Collagen Matrices with Tumor-Mimetic Gradients in Fiber Alignment, Adv Funct Mater 34(13) (2024) 10.1002/adfm.202308071.

[21] H. Zhong, S. Zhou, S. Yin, Y. Qiu, B. Liu, H. Yu, Tumor microenvironment as niche constructed by cancer stem cells: Breaking the ecosystem to combat cancer, J Adv Res (2024) 10.1016/j.jare.2024.06.014.

[22] T. Dou, J. Li, Y. Zhang, W. Pei, B. Zhang, B. Wang, Y. Wang, H. Jia, The cellular composition of the tumor microenvironment is an important marker for predicting therapeutic efficacy in breast cancer, Front Immunol 15 (2024) 1368687, 10.3389/fimmu.2024.1368687.

[23] W. Zhang, M. Wang, C. Ji, X. Liu, B. Gu, T. Dong, Macrophage polarization in the tumor microenvironment: Emerging roles and therapeutic potentials, Biomed Pharmacother 177 (2024) 116930, 10.1016/j.biopha.2024.116930.

[24] M. Sadeghi, S. Dehnavi, M. Sharifat, A.M. Amiri, A. Khodadadi, Innate immune cells: Key players of orchestra in modulating tumor microenvironment (TME), Heliyon 10(5) (2024) e27480, 10.1016/j.heliyon.2024.e27480.

[25] A. Shapouri-Moghaddam, S. Mohammadian, H. Vazini, M. Taghadosi, S.A. Esmaeili, F. Mardani, B. Seifi, A. Mohammadi, J.T. Afshari, A. Sahebkar, Macrophage plasticity, polarization, and function in health and disease, J Cell Physiol 233(9) (2018) 6425–6440, 10.1002/jcp.26429.

[26] M.L. Novak, T.J. Koh, Macrophage phenotypes during tissue repair, J Leukoc Biol 93(6) (2013) 875–881, 10.1189/jlb.1012512.

[27] U. Basak, T. Sarkar, S. Mukherjee, S. Chakraborty, A. Dutta, S. Dutta, D. Nayak, S. Kaushik, T. Das, G. Sa, Tumor-associated macrophages: an effective player of the tumor microenvironment, Front Immunol 14 (2023) 1295257, 10.3389/fimmu.2023.1295257.

[28] P.V. Taufalele, W. Wang, A.J. Simmons, A.N. Southard-Smith, B. Chen, J.D. Greenlee, M.R. King, K.S. Lau, D.C. Hassane, F. Bordeleau, C.A. Reinhart-King, Matrix stiffness enhances cancer-macrophage interactions and M2-like macrophage accumulation in the breast tumor microenvironment, Acta Biomater 163 (2023) 365–377, 10.1016/j.actbio.2022.04.031.

[29] A. Bobrie, O. Massol, J. Ramos, C. Mollevi, E. Lopez-Crapez, N. Bonnefoy, F. Boissiere-Michot, W. Jacot, Association of CD206 Protein Expression with Immune Infiltration and Prognosis in Patients with Triple-Negative Breast Cancer, Cancers (Basel) 14(19) (2022) 10.3390/cancers14194829.

[30] C. Fang, M.Y. Cheung, R.C. Chan, I.K. Poon, C. Lee, C.C. To, J.Y. Tsang, J. Li, G.M. Tse, Prognostic Significance of CD163+ and/or CD206+ Tumor-Associated Macrophages Is Linked to Their Spatial Distribution and Tumor-Infiltrating Lymphocytes in Breast Cancer, Cancers (Basel) 16(11) (2024) 10.3390/cancers16112147.

[31] E. Strack, P.A. Rolfe, A.F. Fink, K. Bankov, T. Schmid, C. Solbach, R. Savai, W. Sha, L. Pradel, S. Hartmann, B. Brune, A. Weigert, Identification of tumor-associated macrophage subsets that are associated with breast cancer prognosis, Clin Transl Med 10(8) (2020) e239, 10.1002/ctm2.239.

[32] J. Sapudom, S. Karaman, W.K.E. Mohamed, A. Garcia-Sabate, B.C. Quartey, J.C.M. Teo, 3D in vitro M2 macrophage model to mimic modulation of tissue repair, NPJ Regen Med 6(1) (2021) 83, 10.1038/s41536-021-00193-5.

[33] R. McWhorter, B. Bonavida, The Role of TAMs in the Regulation of Tumor Cell Resistance to Chemotherapy, Crit Rev Oncog 29(4) (2024) 97–125, 10.1615/CritRevOncog.2024053667.

[34] Y. Heng, X. Zhu, H. Lin, M. Jingyu, X. Ding, L. Tao, L. Lu, CD206(+) tumor-associated macrophages interact with CD4(+) tumor-infiltrating lymphocytes and predict adverse patient outcome in human laryngeal squamous cell carcinoma, J Transl Med 21(1) (2023) 167, 10.1186/s12967-023-03910-4.

[35] W. Liu, H. Zhou, W. Lai, C. Hu, R. Xu, P. Gu, M. Luo, R. Zhang, G. Li, The immunosuppressive landscape in tumor microenvironment, Immunol Res (2024) 10.1007/s12026-024-09483-8.

[36] E.J. Hoffmann, S.M. Ponik, Biomechanical Contributions to Macrophage Activation in the Tumor Microenvironment, Front Oncol 10 (2020) 787, 10.3389/fonc.2020.00787.

[37] M.J. Mitchell, R.K. Jain, R. Langer, Engineering and physical sciences in oncology: challenges and opportunities, Nat Rev Cancer 17(11) (2017) 659–675, 10.1038/nrc.2017.83.

[38] J. Sapudom, T. Pompe, Biomimetic tumor microenvironments based on collagen matrices, Biomater Sci 6(8) (2018) 2009–2024, 10.1039/c8bm00303c.

[39] J. Sapudom, W.K.E. Mohamed, A. Garcia-Sabate, A. Alatoom, S. Karaman, N. Mahtani, J.C. Teo, Collagen Fibril Density Modulates Macrophage Activation and Cellular Functions during Tissue Repair, Bioengineering (Basel) 7(2) (2020) 10.3390/bioengineering7020033.

[40] J. Sapudom, P. Riedl, M. Schricker, K. Kroy, T. Pompe, Physical network regimes of 3D fibrillar collagen networks trigger invasive phenotypes of breast cancer cells, Biomater Adv 163 (2024) 213961, 10.1016/j.bioadv.2024.213961.

[41] M. Nair, R.K. Johal, S.W. Hamaia, S.M. Best, R.E. Cameron, Tunable bioactivity and mechanics of collagen-based tissue engineering constructs: A comparison of EDC-NHS, genipin and TG2 crosslinkers, Biomaterials 254 (2020) 120109, 10.1016/j.biomaterials.2020.120109.

[42] J. Sapudom, L. Kalbitzer, X. Wu, S. Martin, K. Kroy, T. Pompe, Fibril bending stiffness of 3D collagen matrices instructs spreading and clustering of invasive and non-invasive breast cancer cells, Biomaterials 193 (2019) 47–57, 10.1016/j.biomaterials.2018.12.010.

[43] B.C. Quartey, J. Sapudom, M. ElGindi, A. Alatoom, J. Teo, Matrix-Bound Hyaluronan Molecular Weight as a Regulator of Dendritic Cell Immune Potency, Adv Healthc Mater 13(8) (2024) e2303125, 10.1002/adhm.202303125.

[44] J. Sapudom, S. Rubner, S. Martin, S. Thoenes, U. Anderegg, T. Pompe, The interplay of fibronectin functionalization and TGF-beta1 presence on fibroblast proliferation, differentiation and migration in 3D matrices, Biomater Sci 3(9) (2015) 1291–1301, 10.1039/c5bm00140d.

[45] J. Sapudom, S. Rubner, S. Martin, T. Kurth, S. Riedel, C.T. Mierke, T. Pompe, The phenotype of cancer cell invasion controlled by fibril diameter and pore size of 3D collagen networks, Biomaterials 52 (2015) 367–375, 10.1016/j.biomaterials.2015.02.022.

[46] K. Franke, J. Sapudom, L. Kalbitzer, U. Anderegg, T. Pompe, Topologically defined composites of collagen types I and V as in vitro cell culture scaffolds, Acta Biomater 10(6) (2014) 2693–2702, 10.1016/j.actbio.2014.02.036.

[47] S.X. Ge, E.W. Son, R. Yao, iDEP: an integrated web application for differential expression and pathway analysis of RNA-Seq data, BMC Bioinformatics 19(1) (2018) 534, 10.1186/s12859-018-2486-6.

[48] C.D. Paul, P. Mistriotis, K. Konstantopoulos, Cancer cell motility: lessons from migration in confined spaces, Nat Rev Cancer 17(2) (2017) 131–140, 10.1038/nrc.2016.123.

[49] A. Ahmed, I.M. Joshi, S. Larson, M. Mansouri, S. Gholizadeh, Z. Allahyari, F. Forouzandeh, D.A. Borkholder, T.R. Gaborski, V.V. Abhyankar, Microengineered 3D Collagen Gels with Independently Tunable Fiber Anisotropy and Directionality, Adv Mater Technol 6(4) (2021) 2001186–2001186, 10.1002/admt.202001186.

[50] A. Ray, Z.M. Slama, R.K. Morford, S.A. Madden, P.P. Provenzano, Enhanced Directional Migration of Cancer Stem Cells in 3D Aligned Collagen Matrices, Biophys J 112(5) (2017) 1023–1036, 10.1016/j.bpj.2017.01.007.

[51] A. Haque, M. Moriyama, K. Kubota, N. Ishiguro, M. Sakamoto, A. Chinju, K. Mochizuki, T. Sakamoto, N. Kaneko, R. Munemura, T. Maehara, A. Tanaka, J.N. Hayashida, S. Kawano, T. Kiyoshima, S. Nakamura, CD206(+) tumor-associated macrophages promote proliferation and invasion in oral squamous cell carcinoma via EGF production, Sci Rep 9(1) (2019) 14611, 10.1038/s41598-019-51149-1.

[52] J.M. Jaynes, R. Sable, M. Ronzetti, W. Bautista, Z. Knotts, A. Abisoye-Ogunniyan, D. Li, R. Calvo, M. Dashnyam, A. Singh, T. Guerin, J. White, S. Ravichandran, P. Kumar, K. Talsania, V. Chen, A. Ghebremedhin, B. Karanam, A. Bin Salam, R. Amin, T. Odzorig, T. Aiken, V. Nguyen, Y. Bian, J.C. Zarif, A.E. de Groot, M. Mehta, L. Fan, X. Hu, A. Simeonov, N. Pate, M. Abu-Asab, M. Ferrer, N. Southall, C.Y. Ock, Y. Zhao, H. Lopez, S. Kozlov, N. de Val, C.C. Yates, B. Baljinnyam, J. Marugan, U. Rudloff, Mannose receptor (CD206) activation in tumor-associated macrophages enhances adaptive and innate antitumor immune responses, Sci Transl Med 12(530) (2020) 10.1126/scitranslmed.aax6337.

[53] H.B. Abdalla, M.H. Napimoga, A.H. Lopes, A.G. de Macedo Maganin, T.M. Cunha, T.E. Van Dyke, J.T. Clemente Napimoga, Activation of PPAR-gamma induces macrophage polarization and reduces neutrophil migration mediated by heme oxygenase 1, Int Immunopharmacol 84 (2020) 106565, 10.1016/j.intimp.2020.106565.

[54] Q. Yao, J. Liu, Z. Zhang, F. Li, C. Zhang, B. Lai, L. Xiao, N. Wang, Peroxisome proliferator- activated receptor gamma (PPARgamma) induces the gene expression of integrin alpha(V)beta(5) to promote macrophage M2 polarization, J Biol Chem 293(43) (2018) 16572–16582, 10.1074/jbc.RA118.003161.

[55] B. Daniel, G. Nagy, Z. Czimmerer, A. Horvath, D.W. Hammers, I. Cuaranta-Monroy, S. Poliska, P. Tzerpos, Z. Kolostyak, T.T. Hays, A. Patsalos, R. Houtman, S. Sauer, J. Francois-Deleuze, F. Rastinejad, B.L. Balint, H.L. Sweeney, L. Nagy, The Nuclear Receptor PPARgamma Controls Progressive Macrophage Polarization as a Ligand-Insensitive Epigenomic Ratchet of Transcriptional Memory, Immunity 49(4) (2018) 615–626 e616, 10.1016/j.immuni.2018.09.005.

[56] B.H. Cha, S.R. Shin, J. Leijten, Y.C. Li, S. Singh, J.C. Liu, N. Annabi, R. Abdi, M.R. Dokmeci, N.E. Vrana, A.M. Ghaemmaghami, A. Khademhosseini, Integrin-Mediated Interactions Control Macrophage Polarization in 3D Hydrogels, Adv Healthc Mater 6(21) (2017) 1700289–1700289, 10.1002/adhm.201700289.

[57] S.B. Weisser, K.W. McLarren, N. Voglmaier, C.J. van Netten-Thomas, A. Antov, R.A. Flavell, L.M. Sly, Alternative activation of macrophages by IL-4 requires SHIP degradation, Eur J Immunol 41(6) (2011) 1742–1753, 10.1002/eji.201041105.

[58] E. Sahin, S. Haubenwallner, M. Kuttke, I. Kollmann, A. Halfmann, A.M. Dohnal, L. Chen, P. Cheng, B. Hoesel, E. Einwallner, J. Brunner, J.B. Kral, W.C. Schrottmaier, K. Thell, V. Saferding, S. Bluml, G. Schabbauer, Macrophage PTEN regulates expression and secretion of arginase I modulating innate and adaptive immune responses, J Immunol 193(4) (2014) 1717–1727, 10.4049/jimmunol.1302167.

[59] B.H. Cha, S.R. Shin, J. Leijten, Y.C. Li, S. Singh, J.C. Liu, N. Annabi, R. Abdi, M.R. Dokmeci, N.E. Vrana, A.M. Ghaemmaghami, A. Khademhosseini, Integrin-Mediated Interactions Control Macrophage Polarization in 3D Hydrogels, Adv Healthc Mater 6(21) (2017) 10.1002/adhm.201700289.

[60] X. Huang, Y. Li, M. Fu, H.B. Xin, Polarizing Macrophages In Vitro, Methods Mol Biol 1784 (2018) 119–126, 10.1007/978-1-4939-7837-3_12.

[61] S. Zhang, X. Yang, L. Wang, C. Zhang, Interplay between inflammatory tumor microenvironment and cancer stem cells, Oncol Lett 16(1) (2018) 679–686, 10.3892/ol.2018.8716.

[62] F.R. Greten, S.I. Grivennikov, Inflammation and Cancer: Triggers, Mechanisms, and Consequences, Immunity 51(1) (2019) 27-41, 10.1016/j.immuni.2019.06.025.

[63] T. Abdul-Rahman, S. Ghosh, S.M. Badar, A. Nazir, G.B. Bamigbade, N. Aji, P. Roy, H. Kachani, N. Garg, L. Lawal, Z.S.B. Bliss, A.A. Wireko, O. Atallah, F.T. Adebusoye, T. Teslyk, K. Sikora, V. Horbas, The paradoxical role of cytokines and chemokines at the tumor microenvironment: a comprehensive review, Eur J Med Res 29(1) (2024) 124, 10.1186/s40001-024-01711-z.

[64] E. Voronov, D.S. Shouval, Y. Krelin, E. Cagnano, D. Benharroch, Y. Iwakura, C.A. Dinarello, R.N. Apte, IL-1 is required for tumor invasiveness and angiogenesis, Proc Natl Acad Sci U S A 100(5) (2003) 2645–2650, 10.1073/pnas.0437939100.

[65] V. Bigatto, F. De Bacco, E. Casanova, G. Reato, L. Lanzetti, C. Isella, I. Sarotto, P.M. Comoglio, C. Boccaccio, TNF-alpha promotes invasive growth through the MET signaling pathway, Mol Oncol 9(2) (2015) 377–388, 10.1016/j.molonc.2014.09.002.

[66] M. Jung, B. Bonavida, Immune Evasion in Cancer Is Regulated by Tumor-Asociated Macrophages (TAMs): Targeting TAMs, Crit Rev Oncog 29(4) (2024) 1–17, 10.1615/CritRevOncog.2024053096.

[67] S. Xu, Q. Wang, W. Ma, Cytokines and soluble mediators as architects of tumor microenvironment reprogramming in cancer therapy, Cytokine Growth Factor Rev 76 (2024) 12–21, 10.1016/j.cytogfr.2024.02.003.

[68] Z. Mai, Y. Lin, P. Lin, X. Zhao, L. Cui, Modulating extracellular matrix stiffness: a strategic approach to boost cancer immunotherapy, Cell Death Dis 15(5) (2024) 307, 10.1038/s41419-024-06697-4.

